# Host-specific adaptation and fitness trade-off of Barley Yellow Dwarf Viruses suggested by experimental evolution through aphid inoculation on multiple Poaceae species

**DOI:** 10.1101/2025.06.20.660669

**Authors:** Lucie Tamisier, Catherine Colson, François Maclot, Peipei Zhang, Xifeng Wang, Frederic Francis, Denis Baurain, Sébastien Massart

**Affiliations:** Plant Pathology Laboratory, Terra, Gembloux Agro-Bio Tech, Université de Liège, Passage des Déportés, 2, 5030 Gembloux, Belgium; InBioS–PhytoSYSTEMS, Eukaryotic Phylogenomics, University of Liège, Bât. B22, Quartier Vallée 1, Chemin de la Vallée 4, 4000 Liège, Belgium; College of Life Sciences, Langfang Normal University, Langfang 065000, China; State Key Laboratory for Biology of Plant Diseases and Insect Pests, Institute of Plant Protection, Chinese Academy of Agricultural Sciences, 100193 Beijing, China; Functional and Evolutionary Entomology, Terra - Gembloux Agro-Bio Tech, University of Liège, Passage des Déportés, 2, 5030 Gembloux, Belgium

## Abstract

Yellow dwarf viruses are damaging viruses infecting cereals. They are able to infect a wide range of host plants belonging to the *Poaceae* family. The ban of neonicotinoids in Europe has resulted in an increasing disease incidence and triggered the need to better understand their emergence and spread. The ability of a Barley yellow dwarf virus (BYDV) population to adapt to different hosts has never been studied. We performed an experimental evolution of two BYDV species (BYDV-PAS and BYDV-PAV) to study their adaptation to four *Poaceae* species (wheat, oat, two-row barley, and six-row barley). After four months of evolution (4 passages from plant to plant), the replicative fitness of the final viral populations was estimated, and the complete viral genomes were sequenced by high-throughput sequencing in pools of BYDV populations. Wildly divergent evolutionary trajectories were obtained, with stable or increased fitness, up to extinctions of viral populations within and among plant species. To understand these results, the composition of viral populations was analysed in detail using single nucleotide polymorphism (SNP) calling, clustering, and haplotype reconstruction methods. Interestingly, adaptation to oat and barley was mainly explained by a combination of BYDV-PAV haplotypes showing specific mutations. In contrast, adaptation to wheat was mainly explained by a combination of BYDV-PAS haplotypes harbouring specific mutations. Moreover, these local adaptations were associated to an adaptation cost in other hosts for some viral populations. The presence of adaptation costs in controlled but realistic conditions opens the door for evaluating practices such as crop mixtures or rotations on fields, as a means to mitigate the impact of BYDV.

**Author summary:** The use of genetically uniform plant resistant varieties in traditional agriculture creates unique environments that facilitate the rapid emergence of highly virulent pathogen populations. In natural ecosystems, host spatial and temporal heterogeneity help limit the outbreak of epidemics. As a result, disease management strategies such as crop mixtures and rotations have been proposed to reduce the selection pressure exerted on pathogen populations and prevent the emergence of “super-infectious” pathogens. These strategies would be particularly relevant against Barley yellow dwarf virus (BYDV), the virus causing the greatest economic losses on cereals, as insecticides controlling the disease are banned in Europe and few resistance genes are currently available. However, the effectiveness of these strategies against BYDV remains to be demonstrated. By experimentally evolving a natural BYDV population on different cereal species through natural transmission (e.g. vector instead of mechanical inoculation), we showed that different combinations of mutations and haplotypes enable the virus to adapt to different cereal species. Moreover, for some viral populations, the combination promoting adaptation to one host resulted in maladaptation in another host. These host-specific adaptations are key elements in the establishment of crop mixtures and rotations in the field. Our results generated in controlled but realistic conditions demonstrate for the first time that these cultural practices could be effective against these viruses.

## Introduction

Yellow dwarf viruses (YDVs) are the most damaging viruses infecting cereal crops, with significant yield losses regularly occurring in wheat, barley and oats [1]. They constitute a complex of at least 12 virus species, 5 being assigned to the genus *Luteovirus* within the family *Tombusviridae* (BYDV-PAV, BYDV-PAS, BYDV-MAV, BYDV-kerII and BYDV-kerIII), 5 being assigned to the genus *Polerovirus* within the family *Solemoviridae* (CYDV-RPV, CYDV-RPS, MYDV-RMV, MaYMV and ScYLV) and 2 unassigned to any genus (WYDV-GPV and BYDV-SGV). Several novel tentative members have also been identified, such as BYDV-GAV and BYDV-OYV [2]. This virus complex has a worldwide distribution, with a wide host range of over 150 *Poaceae* species and more than 25 aphid species transmitting the viruses in a persistent mode [3,4]. Various symptoms may appear following infection depending on the host plant, including dwarfing of shoots, leaf discoloration, reduction in plant size or reduction in grain size and production.

Control of the YDVs has involved several disease management strategies. Over the past decades, insecticides (primarily pyrethroids and neonicotinoids) have been widely used to control aphid vector populations through seed or vegetation treatment [5]. Numerous studies have highlighted the harmful effects of neonicotinoids, particularly due to their toxicity to non-target pollinating insects [6,7]. These findings led to a progressive ban on their use in the European Union since 2018. In addition, the emergence of insecticide-resistant aphid genotypes also compromises their use [8]. As a result, BYDV is becoming an increasingly important threat and it is urgent to better understand the drivers of the disease emergence and spread in order to design appropriate control strategies. This includes studying plant-virus interactions to develop resistant or tolerant crop varieties.

The search for genetic resistance and tolerance to YDVs in cereals has been the focus of considerable effort, as these genes generally confer effective protection with low environmental impact. Although no effective sources of resistance and tolerance have been found in cultivated cereal species, several resistance and tolerance genes have been identified and introgressed from wild relatives [9]. Variable results have been obtained so far: several varieties have shown weak durable resistance [10] or a restricted spectrum of action, targeting only certain species or strains of the YDV complex [9]. Moreover, the utilization of tolerant plants still poses a potential risk, as it may result in an increase in virus prevalence in the field, thereby heightening the risk of infection for other host plants.

In this regard, cultural practices could play a key role in the control of YDVs. For instance, combining multiple resistant hosts over time (by rotating different cereal varieties or species in the field) or space (by mixing different cereal varieties or species in the field) could help to increase resistance durability through various mechanisms. These mechanisms include the lower availability of a host variety/species and the distance (in space or time) between plants from the same variety/species, which will lead to a dilution effect and reduce pathogen inoculum produced on a specific host species [11]. Moreover, the virus evolutionary potential can be constrained by the presence of adaptation costs to the host. Indeed, if the adaptation cost is strong enough, the different plant hosts in the field could lead to a diversifying selection towards the virus, which will not be able to adapt to every cereal species or variety [12]. Several studies have already reported the efficiency of cereal mixtures in controlling fungal pathogens [13–16]. Some research is starting to be conducted on the effect of intercropping on aphid pressure and YDVs incidence [17,18], but this research is still scarce and more studies are needed. Although promising, variety/species mixing and rotation strategies rely on the strong assumption that adaptation costs exist, so the virus will not be able to adapt to all hosts in diversified natural context. However, the ability of YDVs populations to adapt to multiple *Poaceae* species simultaneously and the molecular basis of this adaptation have never been studied. Moreover, the effectiveness of the mixture/rotation practices with respect to generalist viruses such as YDVs is questionable, especially given its large host range and ability to persist in changing environments [19].

Finally, studying the evolutionary potential of RNA viruses such as YDVs on several host species is a complex task. Indeed, RNA viruses do not exist within their host as a single genome but as a population of related but non-identical genome sequences, exhibiting a high level of genetic diversity. Being able to characterise the whole viral population, including the occurrence of low-frequency resistance-breaking variants, is essential to manage disease-resistant crops in the field. In the past decade, the development of high-throughput sequencing (HTS) technologies has revolutionised the study of viral populations. Thanks to the large numbers of sequences generated by HTS, the viral population, including low-frequency variants, can be characterised with unprecedented resolution [20–23]. For this purpose, several methods have been employed on HTS datasets for exploring SNP diversity [24–26] or for reconstructing haplotypes [27,28], including at national scale [29]. The latter is computationally challenging but makes it possible to determine which SNPs belong to the same viral molecule (*i.e.,* haplotype) in order to better describe the viral population. Therefore, working with haplotypes provides an accurate evaluation of the viral population at the molecular level.

In this study, we performed an experimental evolution of YDVs to investigate their adaptation to four *Poaceae* species. To ensure the most realistic experimental conditions, we initiated the experiment with a composite population consisting of two BYDV species (BYDV-PAS and BYDV-PAV), reflecting the frequent co-infections observed in natural conditions [30–32], and we performed inoculations using aphid vectors. Although mechanical inoculation is the most widely transmission pathway used in experimental evolution, it can introduce a bias in viral populations structure (higher viral diversity) compared to the natural transmission by aphid vectors, as shown for potato virus Y [33]. The replicative fitness of the viral populations was assessed after four months of evolution. In order to interpret the observed evolutionary trajectories, HTS was used to investigate the composition of the viral populations down to haplotype resolution and detect low-frequency variants. With this experiment, we aimed to address three questions: (i) Can different evolutionary trajectories be obtained by evolving a BYDV population on different *Poaceae* species? (ii) Does an accurate description of the viral population compositions, combining mutational analyses, clustering analyses and haplotype reconstruction, can help to understand BYDV evolutionary trajectories? (iii) Can these results allow us to estimate whether agricultural practices such as crop mixtures and rotations could be efficient against generalist viruses like YDVs?

## Results

### From extinction to fitness gain: divergent evolutionary trajectories for BYDV populations, whatever the host species

To study virus adaptation to *Poaceae* species, we conducted an experimental evolution with a mixed population of BYDV-PAV and BYDV-PAS. For four months, four successive passages were carried out through aphid transmission. Fifteen independent viral populations were initiated per host species: wheat (*Triticum aestivum* cv. Avatar) (W), oat (*Avena sativa* cv. Evita) (O), two-row barley (*Hordeum vulgare* cv. Laureate) (TB), and six-row barley (*Hordeum hexastichum* cv. Smooth) (SB) (Fig 1).

**Figure 1:**
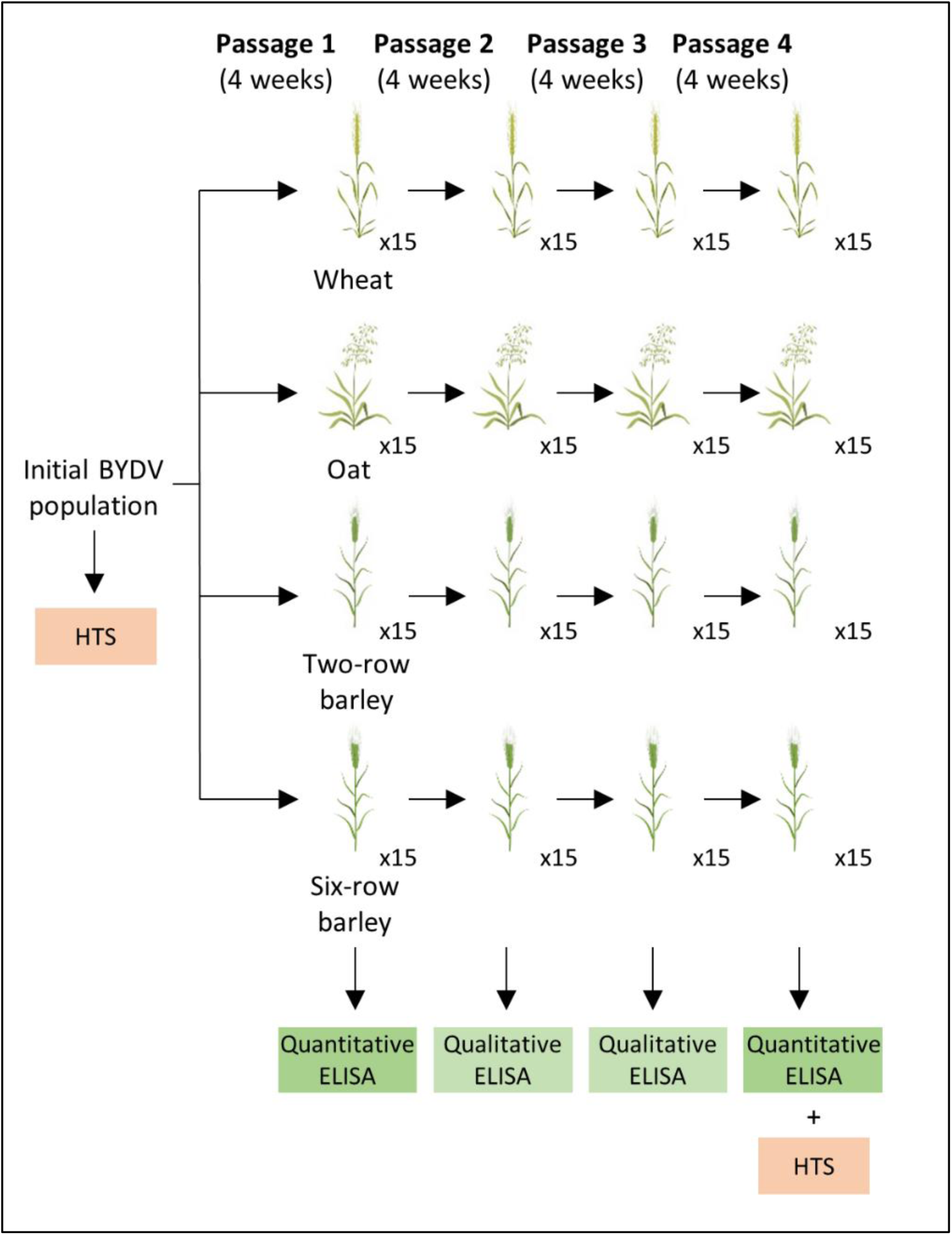
Experimental design of the study. Initial BYDV population was inoculated in four *Poaceae* species, with fifteen plants per species. Viral populations were passed every month in the same species for four months, using five *Rhopalosiphum padi* aphids per plant to perform the inoculation. The initial BYDV population and pool of plants infected with the final BYDV populations were sequenced using high-throughput sequencing (HTS) technology. A quantitative (passages 1 and 4) or qualitative (passages 2 and 3) Enzyme-Linked Immunosorbent Assay (ELISA) was performed on each individual plant.

The mean viral accumulation of the BYDV populations significantly increased in all host species between the 1^st^ and 4^th^ passages (Wilcoxon test, p < 0.001). A similar pattern was observed for the four host species with three contrasting situations among individual plants: significantly increased viral accumulation (4 to 8 plants *per* host), no change (2 to 6 plants), or decrease/extinction (1 to 8 plants) (Fig 2). The number of extinctions varied between 0 and 4 after a single passage depending on the host species, but on average was 10% at each passage (S1 Table). The increase in viral accumulation varied greatly among viral populations evolving in the same host species, for example ranging from 3.6-fold (population P6) to 139-fold (population P1) in oat. Within-host viral accumulation is a component of viral fitness, which is called replicative fitness [34], and will be referred to as fitness in the rest of the article.

**Figure 2:**
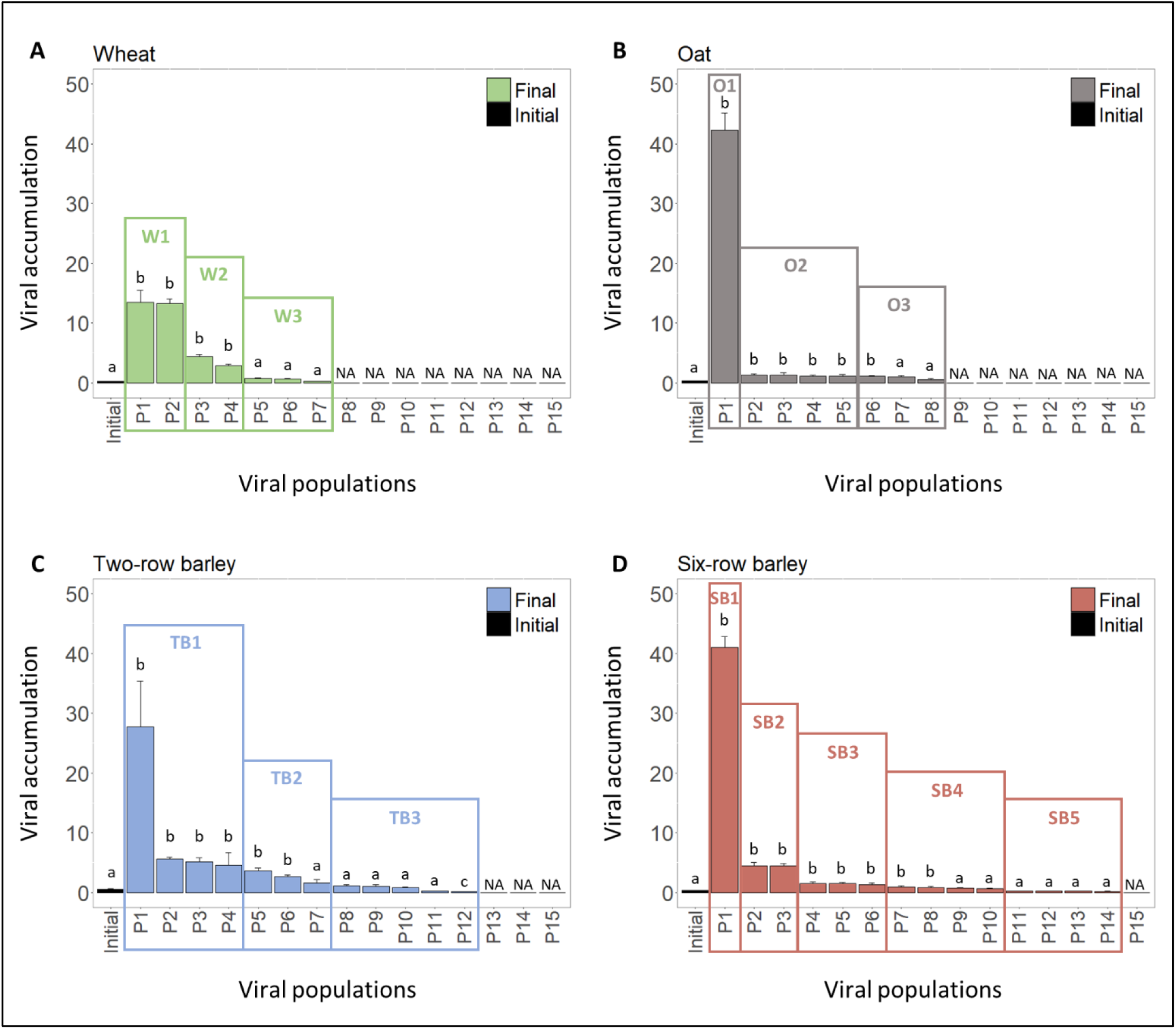
Evolution of the viral accumulation over the experiment. Average viral accumulations of the initial BYDV population after the first passage (initial in black) and individual viral accumulations of the 15 final BYDV populations after four months of evolution on wheat (A), oat (B), two-row barley (C), and six-row barley (D) are represented. For each host species, initial viral accumulation was calculated as the average of the viral accumulation measured in 15 individual plants after the 1^st^ passage. Final viral accumulation within each plant was calculated as the average of the viral accumulation measured four times after grinding of the plant. Units are expressed as relative concentration, comparing the viral accumulation of each plant to a control reference. Dunnett t-testing was conducted to check for differences between the initial BYDV population and the final BYDV populations in each plant (P1 to P15). Letters a, b and c indicate differences among the groups obtained after the test, the final populations showing no differences (letter a), a significant higher viral accumulation (letter b) or a significant lower viral accumulation (letter c) compared to the initial population, respectively. Viral populations pooled to perform the sequencing are surrounded and pool names are indicated (W1 to W3 for wheat; O1 to O3 for oat; TB1 to TB3 for two-row barley; SB1 to SB5 for six-row barley). NA: not measured (extinct lineage). Error bars indicate standard error of the mean. The 15 viral populations have been sorted by viral accumulation level.

### The mutational landscape is dominated by unique mutations with few parallel mutations within and between host species

To investigate the genetic origin of these fitness differences, high-throughput sequencing (HTS) was performed on the initial BYDV population and on plants of the 4^th^ passage pooled according to the viral accumulation of individual plants: three pools for wheat, oat and two-row barley, and five pools for six-row barley (Fig 2). The genetic diversity of the pooled BYDV populations was analysed using three approaches: (i) mutational analyses, (ii) clustering analyses, and (iii) haplotype reconstruction analyses.

The sequencing of the initial BYDV population generated two nearly complete genomes of 5,577 and 5,570 nt (GenBank accessions OM046619 and OM046620) corresponding to BYDV-PAS and BYDV-PAV, respectively. The taxonomic assignation was confirmed by pairwise distance analyses (S2 Table). The genomes showed 87% identity at the nucleotide level. BYDV-PAV was the dominant species in the initial sample with a frequency of 87%. Both BYDV genomes were used as reference genomes for mapping individual reads and single nucleotide polymorphism (SNP) analyses. The average genome sequencing depth for all the samples ranged between 1,116× and 26,443× (S3 Table). After four months of evolution, 136 and 92 genomic positions presented mutated bases with a frequency higher than 1% for BYDV-PAS and BYDV-PAV, respectively (Fig 3). Among these genomic positions in the 14 sequenced pooled populations, a total of 667 and 146 SNP events were detected. *De novo* mutations (*i.e,* mutations absent from the initial BYDV population) were detected at 86 (109 SNP events) and 82 (96 SNP events) genomic positions for BYDV-PAS and BYDV-PAV, respectively. For both species, 78% of the *de novo* mutations showed a relatively low frequency (1 - 10%) and around 85% of the *de novo* mutations were present in a single sequenced population (Fig 3, Table 1). In addition, parallel *de novo* mutations (*i.e.,* a mutation appearing at the same genomic position in independently evolving population) were observed in several plant species (n=12 and 4 for BYDV-PAS and BYDV-PAV, respectively) or between different populations in a single plant species (n=2 and 7 for BYDV-PAS and BYDV-PAV, respectively).

**Figure 3:**
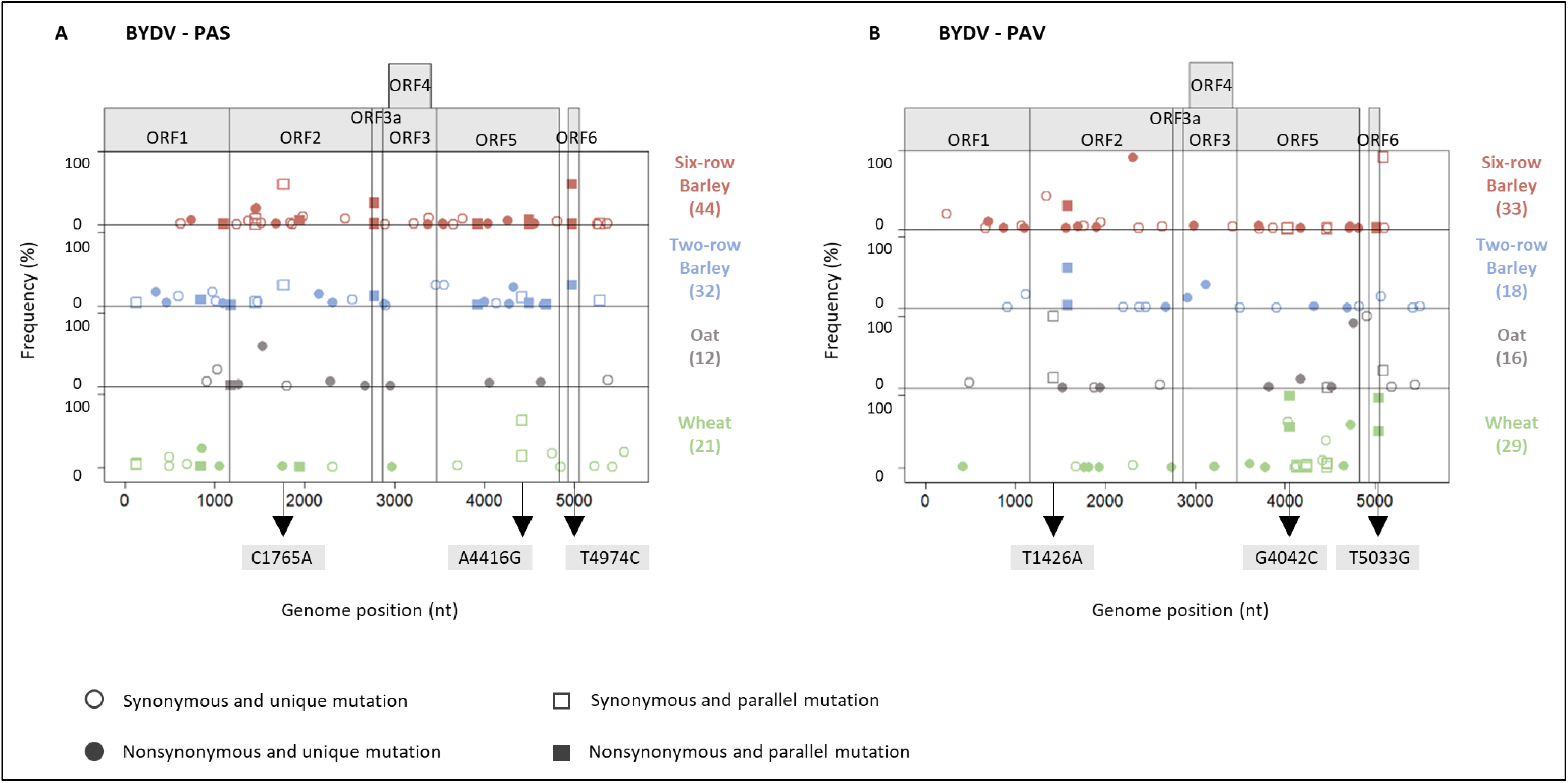
Representation of the frequencies and positions of the *de novo* mutations identified in the final BYDV pooled populations. Frequencies of *de* mutations detected are plotted according to their positions in BYDV-PAS (A) and BYDV-PAV (B) genomes. Symbol colours indicate the host species in the mutations were identified: wheat (green), oat (grey), two-row barley (blue), or six-row barley (red). Total number of *de novo* mutations in each host es is given in brackets. Unique and parallel mutations are represented by a circle and a square, respectively. Synonymous and nonsynonymous mutations presented by an empty and a filled symbol, respectively. Parallel *de novo* mutations mostly carried by viral pooled populations showing increased in fitness ith a frequency higher than 10% in at least one pooled population are indicated in grey boxes.

**Table 1:** Summary statistics of the *de novo* mutations generated during the experimental evolution.

| Mutations | BYDV-PAS | BYDV-PAV |
| --- | --- | --- |
| Total | 86 | 82 |
| Total synonymous | 45 | 44 |
| Total nonsynonymous | 41 | 38 |
| Proportion of rare mutations (1%-10%) | 78% | 78% |
| Mutations unique to a viral population | 72 | 71 |
| Parallel mutations appearing in at least two plant species | 12 | 4 |
| Parallel mutations appearing only in the same plant species | 2 | 7 |

For BYDV-PAS, three parallel mutations were found at relatively high frequencies and mostly appeared in viral populations showing increased fitness (Fig 3A, Fig 2). The mutation C1765A (synonymous and in ORF2 coding for RNA-dependent RNA polymerase (RdRp)) was shared by SB1 and TB2 at frequencies of 56% and 28%, respectively. The mutation A4416G (synonymous and in ORF5 which encodes a CP-readthrough fusion protein, which is necessary for transmission via aphids) was present in W1, W2, and TB3 at 64%, 16% and 12% frequencies, respectively. The mutation T4974C (nonsynonymous, valine to alanine, and in ORF6, encoding a small protein with no function assigned) was shared by SB1, TB2, and SB4, at 55%, 28% and 1% frequencies, respectively.

For BYDV-PAV, three parallel mutations showed high frequencies and were detected in viral populations showing increased fitness (Fig 3B, Fig 2). Mutation T1426A (synonymous and in ORF2 coding for RdRp) was observed in O1 and O2, at 99.9% and 14.9%, respectively. In addition, W1 and W2 shared two specific mutations at similar frequencies (96-99% in W1 and 50-55% in W2): G4042C (nonsynonymous, glutamic acid to glutamine, and in ORF5) and T5033G (nonsynonymous, serine to alanine, and in ORF6). Therefore, these mutations could be localised on the same haplotype.

### The evolution of population structure is independent between co-infecting viral species

A recently published methodology based on SNP patterns, fixation index (FST) calculation and Principal Component Analysis (PCA) was applied to explore the population structure of both viral species beyond the consensus genomes. This methodology improves the resolution of population structure and can identify associations between SNPs [29]. All the pre-existing and *de novo* SNPs detected in the viral pooled populations, along with their relative frequencies, were computed through an FST approach. Dendrograms were constructed from pairwise distance matrices, and approximately unbiased (AU) p-values were calculated. The dendrogram obtained from the detected SNPs for both species were presented (Fig 4A and Fig 5A). Analysis of the clusters showed that each BYDV species had followed its evolutionary trajectory independently. Indeed, none of the branches of a BYDV species in the dendrogram was reproduced for the other viral species. In addition, the grouping of the viral populations in the PCA was very different between both species (Fig S1). Then, the in-depth analysis of viral populations was carried out separately for each species.

**Figure 4:**
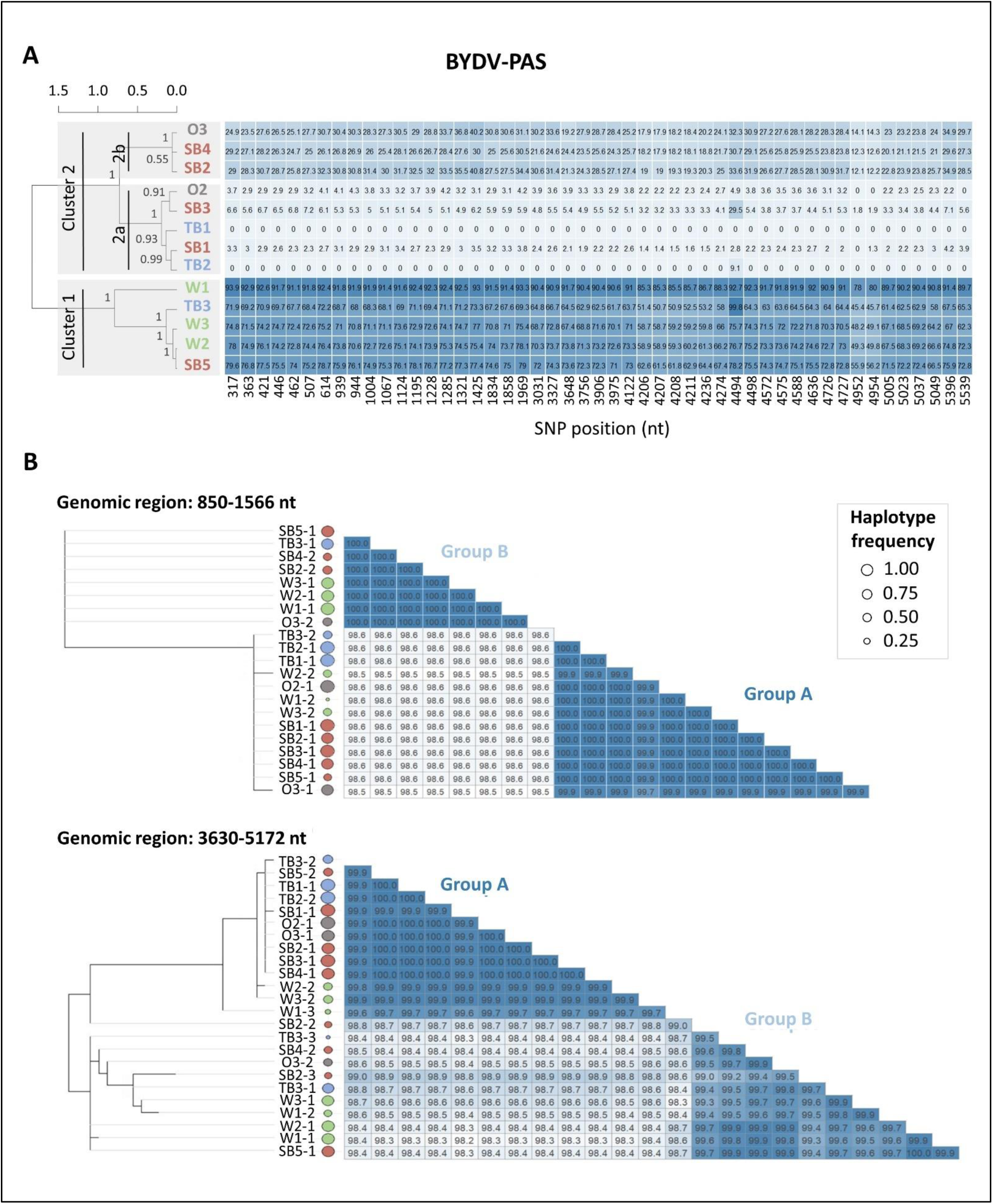
Clustering analysis and phylogenetic trees of BYDV-PAS. Dendrograms show the hierarchical clustering of the final BYDV-PAS pooled populations (A). Dendrograms were constructed from pairwise FST matrices based on all the SNPs detected in the BYDV-PAS pooled populations. Names of the viral populations are indicated (W1 to W3 for wheat; O1 to O3 for oat; TB1 to TB3 for two-row barley; SB1 to SB5 for six-row barley). Numbers indicate the approximately unbiased (AU) p-values. The table shows the frequencies of 49 SNPs contributing to more than 90% of the first component of the PCA. BYDV-PAS haplotypes composing the viral pooled populations were reconstructed using two high SNP density genomic regions: 850-1566 nt and 3630-5172 nt (B). Unrooted phylogenetic trees were obtained using maximum likelihood under the HKY85 model of sequence evolution and are drawn for the BYDV-PAS haplotypes reconstructed for each genomic region. Percentage identity matrices for the nucleotide alignment of the haplotypes are plotted next to the phylogenetic trees. Frequency of each haplotype within the viral population is proportional to circle area. Colour of the circle indicates the host species in which the haplotypes have evolved: wheat (green), oat (grey), two-row barley (blue) or six-row barley (red). The haplotype number is given following the name of the viral population.

**Figure 5:**
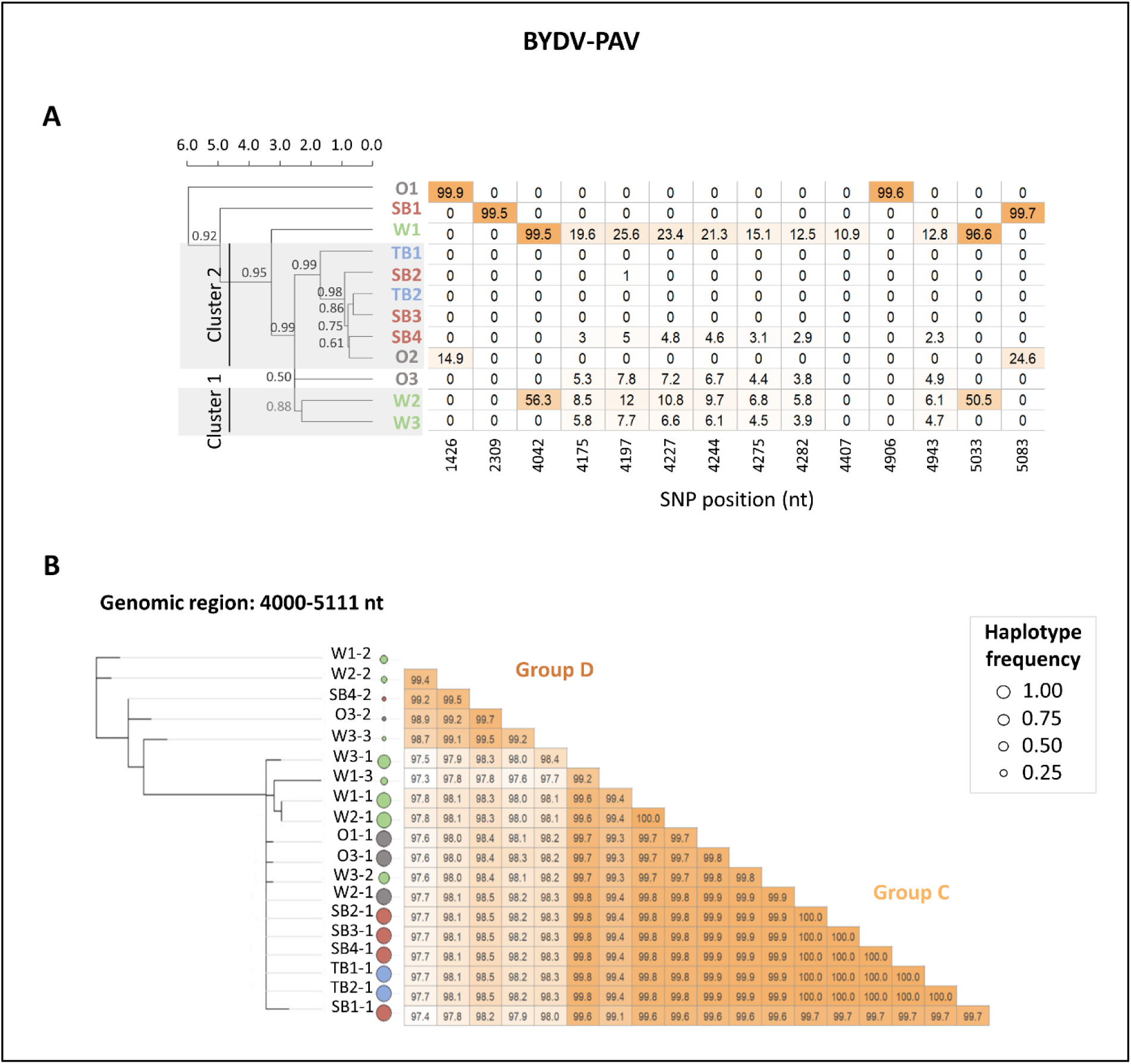
Clustering analysis and phylogenetic trees of BYDV-PAV. Dendrograms show the hierarchical clustering of the final BYDV-PAV pooled populations (A). Dendrograms were constructed from pairwise FST matrices based on all the SNPs detected in the BYDV-PAV pooled populations. Names of the viral populations are indicated (W1 to W3 for wheat; O1 to O3 for oat; TB1 to TB3 for two-row barley; SB1 to SB5 for six-row barley). Numbers indicate the approximately unbiased (AU) p-values. The table shows the frequencies of 14 SNPs contributing to more than 90% of the first component of the PCA. BYDV-PAV haplotypes composing the viral pooled populations were reconstructed using one high SNP density genomic region (B). Unrooted phylogenetic trees were obtained using maximum likelihood under the HKY85 model of sequence evolution and are drawn for the BYDV-PAV haplotypes reconstructed in the genomic region 4000 – 5111 nt. Percentage identity matrices for the nucleotide alignment of the haplotypes are plotted next to the phylogenetic trees. Frequency of each haplotype within the viral population is proportional to circle area. Colour of the circle indicates the host species in which the haplotypes have evolved: wheat (green), oat (grey), two-row barley (blue) or six-row barley (red). The haplotype number is given following the name of the viral population.

### Haplotype reconstruction to decipher the complexity of population structure of BYDV-PAS

For BYDV-PAS, two main clusters were found with a high distance between them and AU p-values of 1 for each cluster (Fig 4A and S1A). The three populations that evolved on wheat belonged to cluster 1, together with TB3 and SB5, the samples with the lowest virus concentration on two-row and six-row barley. Cluster 2 was separated into two sub-clusters (2a and 2b with AU p-values of 1), each including a mix of host species. These results were confirmed by performing a principal component analysis (PCA) using the frequency of the 667 SNPs as variables (Fig S1A). The first component explained 41.6% of the variance and grouped the pooled populations in the same three clusters as the dendrogram. The viral populations were not clustered according to the host species but this PCA did not take into account the fitness of the population nor the haplotype composition.

Forty-nine SNPs, contributing to 90% of the first component of the PCA, were extracted and their frequencies were analysed. The 49 SNPs, distributed along the whole genome, were either always present at similar frequencies, or absent from the viral pooled populations (Fig 4A). Moreover, all these SNP were already pre-existing in the initial BYDV population. The SNP frequencies explained the clusters obtained with the dendrograms. The viral population from cluster 2a presented SNP frequencies ranging between 0 and 7% (excepting a frequency of 27% for SNP at position 4494 for SB3), while the SNP frequencies of cluster 2b ranged from 12 to 41%. Cluster 1 presented high SNP frequencies (between 45.0 and 99.8%). These results suggested that these SNPs belong to the same molecule and constitute a haplotype. This implies that at least two major BYDV-PAS haplotypes were present in the initial inoculum at the beginning of the experiment and that their differences in frequency explained the clustering of the viral pooled populations in the dendrogram.

Determining the sequence of the major and minor haplotypes would allow a refined understanding of the genetic structure of a viral population. Therefore, the haplotype sequences of BYDV-PAS were reconstructed on the sole high SNP density regions to obtain robust results and avoid chimeric haplotypes. Two genomic regions were selected for BYDV-PAS: 850 - 1566 nt (region 1) and 3630 - 5172 nt (region 2), which included 12 and 28 SNPs previously selected for their contribution to the discrimination of the viral pooled populations, respectively. Twenty-one and twenty-four haplotype sequences were reconstructed for regions 1 and 2, respectively, with a maximum of three different haplotypes per viral population (S4 Table). Unrooted phylogenetic trees and pairwise distance analysis revealed two main haplotype groups (A and B) consistent with the whole-genome SNP analysis (Fig 4B). Based on their most frequent haplotype sequence, the samples clustered in the same groups compared to the SNP whole genome analysis. Haplotype sequences showed high identity within groups (>99.9% in region 1) but diverged by up to 1.7% between groups. The sequence of the viral molecules could therefore be reconstructed despite their very high identity, but only on portion of the genomes with high SNP density. For both genomic regions, the samples clustered identically based on their most frequent haplotype sequence, suggesting that the sequences clustering in group A or in group B for each region came from the same viral molecule and belonged to the same haplotype. The two haplotype groups A and B were also found in the initial population, with frequencies of around 96% for group A and 4% for group B.

### Haplotype reconstruction to decipher the complexity of population structure of BYDV-PAV

The same analysis was carried out for BYDV-PAV. In the dendrogram, based on 146 SNPs, populations showing high fitness, such as O1, SB1, and W1 did not cluster with any other populations, while the other pooled populations composed a cluster with AU p-values of 0.99 (Fig 5A). This cluster contained two sub-clusters: W2 and W3 (cluster 1) with AU p-values of 0.88; and TB1, SB2, TB2, SB3, SB4, and O2 (cluster 2) with AU p-values of 0.99.

Again, the PCA results were relatively consistent with those of the dendrogram, with the first component explaining 22% of the variance and grouping the viral pooled populations in three groups: W1; W2 and W3; and the remaining populations (Fig S1B). Fourteen SNPs contributing to 91.8% of the first component were extracted and their frequencies analysed (Fig 5A). The SNPs pattern was very different from BYDV-PAS: eight SNPs (between positions 4175 and 4943) presented similar frequencies in several viral pooled populations and most probably belonged to the same molecule. These 8 SNPs were already pre-existing in the initial BYDV population. The other SNPs are *de novo* SNPs and were absent in most of the samples but present at high frequency (10.9% to 99.9%) in a single (n=2) or two (n=4) populations. Interestingly, the three populations (W1, SB1 and O1) that did not cluster with any other pooled population and showed the highest increase in viral accumulation (Fig 2) presented specific SNP patterns. For instance, populations SB1 and O1 only had two *de novo* SNPs each, but these SNPs were almost fixed in the populations (positions 1426 and 4906 nt for O1; positions 2309 and 5083 nt for SB1). The other almost fixed *de novo* SNPs were previously presented (Fig 3).

To reconstruct the haplotype sequences, a region spanning 1112 nt with a high SNP density (n=12) was selected (positions 4000 - 5111). In total, 19 haplotype sequences were reconstructed, with a maximum of 3 haplotype sequences for a sample (Fig 5B). Two main groups, called C and D, were determined. Group D included all haplotypes carrying the group of eight previously identified SNPs present at a low frequency in all viral populations that evolved on wheat (W1, W2, and W3), as well as in O3 and SB4, but absent from other populations. BYDV-PAV haplotypes from group C were therefore always dominant. The two haplotype groups C and D were also found in the initial population, with frequencies of around 99% for group C and 1% for group D.

### The host species impacted the composition and the concentration of the viral populations at haplotype level

After characterizing the final viral pooled populations at the haplotype level, the link between their haplotype composition, the presence of *de novo* mutations, and viral accumulation was examined (Fig 6). The composition of the final viral pooled populations evolved in a similar way in oat, two-row barley and six-row barley. In barley, the viral pooled populations that least accumulated (TB3 and SB5) were only composed of BYDV-PAS, with more than 70% of the populations composed of BYDV-PAS from group B. In oat, the viral pooled population that least accumulated (O3) was composed of BYDV-PAS from both groups as well as BYDV-PAV from group C. As viral accumulation increased, the proportion of BYDV-PAS in the viral pooled populations tended to decrease in these three host species. Additionally, BYDV-PAS from group B disappeared frequently. The only exception is SB1, which shows an extremely high viral accumulation level with 48% BYDV-PAS (considered as outlier and not included in the linear regression in Figure 6B).

**Figure 6:**
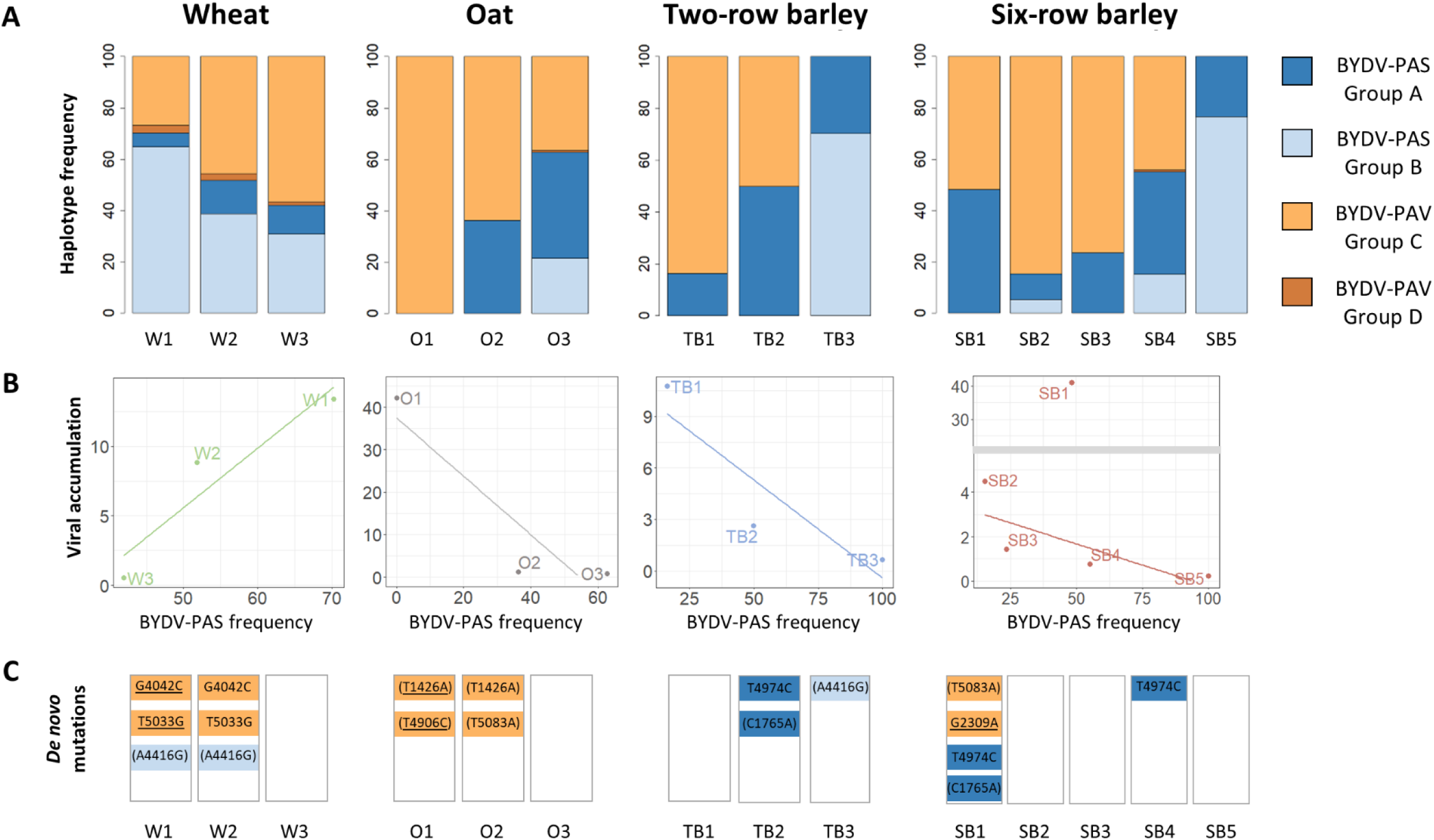
Haplotype composition of the viral pooled populations. Bar plots show the frequencies of the four main haplotype groups in each viral pooled population for BYDV-PAS group A (dark blue), BYDV-PAS group B (light blue), BYDV-PAV group C (light orange) and BYDV-PAV group D (dark orange) (A). For each host species, viral accumulation measured after 4 months of evolution is shown as a function of the BYDV-PAS frequency in the final populations (B). Linear regressions illustrate the trend in viral accumulation relative to the proportion of BYDV-PAS. SB1 was considered an outlier and was not included in the linear regression. Parallel and/or fixed *de novo* mutations mostly carried by viral pooled populations showing increased fitness are represented within each viral population and haplotype group (C). Colour of the mutations indicates the haplotype group in which they appeared. A mutation name between brackets corresponds to a synonymous mutation, while a mutation name without brackets corresponds to a nonsynonymous mutation. An underlined mutation name indicates a fixed mutation in the viral population.

Several mutations, whether fixed and/or shared by multiple viral populations, could also explain the evolution of the viral populations, especially in O1 and SB1, which both showed an extremely high level of viral accumulation (> 40). In O1, the two mutations were identified in BYDV-PAV from group C and were fixed and synonymous. As presented before, the T1426A mutation was also shared by O2. In SB1, mutation G2309A in BYDV-PAV was nonsynonymous and localised in the RdRp region. SB1 also harboured mutation T4974C in BYDV-PAS, which was shared with TB2 and SB4, as well as mutation C1765A shared with TB2.

The viral pooled populations that evolved on wheat were differently composed from the other viral pooled populations (Fig 6A). The most prevalent haplotypes on wheat were BYDV-PAS from group B and BYDV-PAV from group C. Moreover, the frequency of BYDV-PAS from group B increased with the rise in viral accumulation, in contrast to what we observed in the other three species (Fig 6B, Fig S1C). BYDV-PAS from group A and BYDV-PAV from group D were also found in these three viral pooled populations, but to a lower extent. Interestingly, the three same mutations were detected in W1 and W2, and these two viral pooled populations accumulated most in wheat (Fig 6C).

### Transfer to another host species suggests the existence of fitness trade-offs

The results obtained from viral pooled populations suggest a fitness trade-off, with some viral haplotypes and populations better adapted to wheat and others to barley, two-row barley, and six-row barley. To test this hypothesis, an additional experiment was conducted using individual (*i.e.*, non-pooled) viral populations to assess their fitness on different host species. Four viral populations, which differed in their fitness levels on the host species they evolved on, were selected: O1 (population P1 – high accumulation), O3 (population P7 – low accumulation), SB1 (population P1 - high accumulation) and SB5 (population P11 – low accumulation) (Fig 2). These populations were then passed on three host species: wheat, oat and six-row barley. The viral accumulation measured one month post inoculation confirmed the previous results: O1 (P1) and SB1 (P1), which evolved and reached high accumulation on oat and six-row barley, respectively, showed significantly better adaptation to their respective host species compared to wheat (Fig 7). In contrast, O3 (P7) and SB5 (P11) having evolved at low accumulation rate on oat and six-row barley, respectively, were significantly better adapted to wheat compared to their previous respective host.

**Figure 7:**
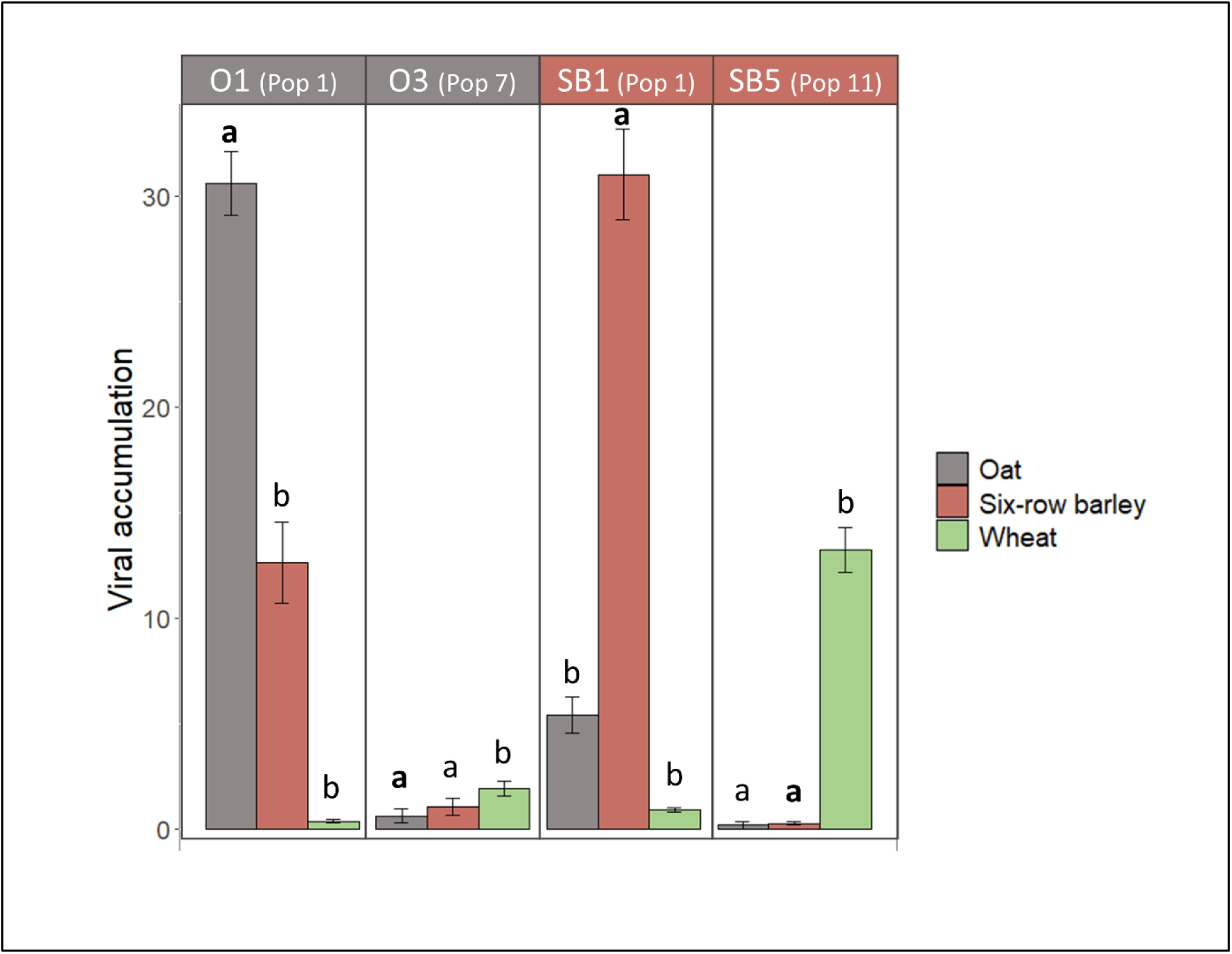
Viral accumulation of 4 viral populations on 3 host species. Viral populations O1 (population P1), O3 (population P7), SB1 (population P1) and SB5 (population P11) were passed on 4 plant species (oat, six-row barley and wheat) with 4 plants per species. Four weeks post inoculation, the viral accumulation in each plant was measured by quantitative ELISA. For each viral population, Dunnett t-testing was used to check for differences between the accumulation of the viral population within the plant species on which it has evolved, and the viral accumulation within the two other plant species. Letters a and b indicate differences in viral accumulation, with no difference with the initial host plant (a) or a significant different with the initial host plant (b). For each viral population, the letter “a” of the control group is indicated in bold.

## Discussion

*Yellow dwarf viruses* (YDVs) are the most important viruses infecting cereals, resulting in substantial economic losses. With the ban of insecticides, the control of these viruses will require the use of genetically resistant or tolerant varieties, as well as a reasoned deployment of these varieties in the field to control the disease over the long term. Crop mixtures and rotations are promising strategies to minimize the adaptive ability of the pathogen. These strategies rely on the action of disruptive selection, which assumes that the viruses will not be able to adapt simultaneously to all the crops deployed. However, the relevance of such practices for generalist viruses such as YDVs, which can adapt to numerous host species, has never been tested.

Understanding how generalist viruses such as YDVs adapt to different host species is therefore crucial for developing sustainable disease management strategies. Our experimental evolution study shows that evolving a composite BYDV population on different *Poaceae* species (wheat, oat, two row barley and six rows barley) using natural transmission pathway (aphids) leads to contrasting evolutionary trajectories. These results provide valuable insights into the mechanisms of viral adaptation and the potential consequences for field-scale control methods. By examining viral population compositions, identifying potential adaptive mutations, and assessing fitness trade-offs, we highlight key factors that could influence the evolution of YDVs in crops.

### Strong bottlenecks induce contrasting evolutionary trajectories within a host species

The initial BYDV population followed markedly divergent evolutionary trajectories when passed to several plants belonging to the same host species. The viral accumulation measured in each individual plant can be viewed as a proxy of the viral population replicative fitness. In each host species, some BYDV populations increased in fitness and adapted to their host, while others did not show any fitness change or became extinct (Fig 2). These different evolutionary trajectories have been observed on viral populations that were initially closely related. The differential adaptation observed among populations could reflect genetic drift, which is a stochastic process leading to the fixation or the loss of mutations independently of their fitness effects. Genetic drift may arise at every step of the infection process, including inoculation, cell to cell movement and host colonisation [35]. During the inoculation process, only a small fraction of the initial viral population composes the inoculum and can establish the new viral population. This strong bottleneck contributes to increase genetic drift. The transmission mode can significantly affect the bottleneck size and the within-plant diversity of the viral population, as demonstrated for *Potato virus Y* [33]. In our study, inoculation was not performed mechanically but by aphid vectors, in a persistent circulative mode, aligning our approach more closely with natural conditions. Several studies have shown that the number of viral particles transmitted *per* aphid is very low: 1 or 2 founders on average are sufficient to start an infection process, for both circulative and non-circulative viruses [36–39]. We chose to use five *Rhopalosiphum padi* aphids per plant for inoculation, as preliminary experiments indicated that this method achieved nearly 100% transmission from wheat to wheat (data not shown). However, this choice likely imposed a significant bottleneck on the viral populations, which may have contributed to the divergent evolutionary trajectories observed, resulting in the elimination of adaptive viral mutations and/or fixation of maladaptive viral mutations in certain populations. While genetic drift could account for some variability, it is noteworthy that certain populations increased their viral accumulation by more than 100-fold between the start and end of the experiment. This suggests that selection also played a crucial role, favoring viral variants that were better adapted to their hosts. These contrasting results allowed the study of the genetic origin of these differences.

### Distinct molecular pathways contribute to local adaptation in BYDV-PAS and BYDV-PAV

In each host species, local adaptations were observed for several viral populations, as it is often observed in evolution experiments [40]. Indeed, numerous studies have shown that when a viral population evolves on a single host, local adaptation occurs, accompanied by the fixation of adaptive mutations [41–49]. We showed that for both BYDV species, local adaptation was due to the selection of both pre-existing viral haplotypes from the initial population and the appearance of *de novo* mutations potentially adaptative (Fig 6). The number of viral mutations that enable host adaptation is often limited [40]. Indeed, the small genome of RNA viruses, with overlapping genes and multifunctional proteins, restricts the number of potential adaptative solutions. Therefore, parallel mutations frequently occur in independent evolutionary populations. Interestingly, our results are partially consistent with these observations. On one hand, several parallel mutations have emerged as candidates for adaptation within each viral species (Mutations C1765A, A4416G and T4974C for BYDV-PAS; Mutations T1426A, G4042C and T5033G for BYDV-PAV) (Fig 3). On the other hand, no potentially adaptive mutations were common between BYDV-PAS and BYDV-PAV, which indicated that within this complex of viral species, the genotypic fitness landscape is not as limited as might be expected. Beyond the viral species, none of the candidate mutations for adaptation was common to the different BYDV haplotypes.

These results could be explained by epistatic interactions between viral mutations, which occur when the fitness value of a mutation depends on the genetic background [50]. This phenomenon has already been described for several plant viruses like the *Tobacco etch virus* [51] or the *Potato virus Y*, where the ability of the virus to breakdown a major pepper resistance gene through mutations in the VPg was impacted by the viral genetic background [52]. In our case, several elements suggested the existence of these epistatic interactions: (i) the consistent emergence of potentially adaptive *de novo* mutations only in a single viral genetic background, and (ii) the independent emergence and subsequent selection of two parallel mutations, G4042C and T5033G, in both the W1 and W2 viral populations, suggesting an epistatic interaction between these two mutations.

Since the effect of a mutation can be affected by the genetic background in which that mutation appears, being able to reconstruct viral haplotypes is crucial to understand viral evolution. High-throughput sequencing has made it possible to study the genetic basis of the evolutionary process, by detecting all mutations present within a population, even at low frequencies. Nevertheless, very few experimental evolution studies have performed haplotype reconstruction so far, due to the difficulty of determining which mutations are physically linked using short reads as well as the frequent errors made by haplotype callers, especially in the presence of high genetic diversity [27]. Using CliqueSNV on targeted genomic region showing high SNP density (*i.e.* two SNPs being no more distant than 150 bp, which allows them to be localised on a single read) [53], we were able to overcome this limitation and reconstruct some of the haplotypes present in our viral populations. This method could miss some haplotypes since the entire viral genome is not utilised, but it prevents the generation of chimeric haplotypes. Haplotype reconstruction has been performed on two distinct genomic regions for BYDV-PAS and similar results have been obtained, two main haplotype groups being detected every time, including at low frequency in the initial population, before the experiment, demonstrating the robustness of this method.

Our method is very well suited for the short-read sequencing technologies, corresponding to the vast majority of generated datasets up to now but, in the coming years, long-read sequencing technologies like Oxford Nanopore MinION or Pacific Biosciences will open up new possibilities and could be an alternative to haplotype reconstruction. Indeed, these technologies can generate read lengths of 10 kb or more and could allow the sequencing of complete haplotype genomes [54].

### Local adaptation is likely associated with a fitness trade-off

Our study also demonstrated that local adaptation to a host was associated, at least for some viral populations, with an adaptation cost: viral populations with high fitness in a first host have significantly lower fitness in a second host (Fig 7). This result could be due to a phenomenon of antagonistic pleiotropy, whereby mutations beneficial on the first host are deleterious on the second host. Although local adaptation is not always associated with fitness trade-offs, numerous examples exist for plant viruses [55].

In our case, antagonistic pleiotropy could occur at two levels. At haplotype level, BYDV-PAS from group B is the most frequent haplotype in the viral populations most adapted to wheat, but also in the populations least adapted to two-row and six-row barley (Fig 6, Fig S1C). Conversely, BYDV-PAV from group C is the most frequent haplotype in the viral populations most adapted to oat, two-row and six-row barley, but also in the populations least adapted to wheat. In addition, cross-inoculation tests confirmed that viral populations composed of BYDV-PAV from group C such as O1 (population P1) had high fitness on oat but significantly lower fitness on wheat, while other populations mainly composed of BYDV-PAS from group B such as SB5 (population P11) had higher fitness on wheat than on two-row barley and six-row barley, the latter being the species on which it evolved during the experiment (Fig 7). As a result, certain haplotype-defining mutations could have antagonistic pleiotropic effects. Antagonistic pleiotropy could also occur at the scale of *de novo* mutations. In our study, populations O1 (population P1) and SB1 (population P1) carry several *de novo* mutations, which could explain their very high level of adaptation on the hosts on which they evolved, as well as their low fitness level on the other two hosts tested (Fig 7). These hypotheses could only be formally validated by testing the effect of each specific mutation on virus fitness in the different viral haplotypes. This would require performing directed mutagenesis using infectious BYDV clones.

### Prospects for implementing BYDV control methods

Our results are encouraging for the implementation of YDVs field-scale control methods, such as varietal or species mixtures and rotations. Indeed, these methods are based on the condition that adaptation costs exist for the pathogen. We showed that, despite the fact that YDVs are generalist viruses, adaptation costs can be identified on different host species. The controlled evolution experiment was performed under the most realistic possible conditions. Firstly, inoculations were carried out using aphid vectors of the virus, as in the field (compared to mechanical inoculation usually carried out in such experiments). Secondly, a mixture of BYDV species and haplotypes (not a single infectious clone as it is often the case in the literature) was used to start the experimental evolution. Indeed, YDVs are a complex of species frequently observed in co-infection in the field [1]. Nevertheless, the transfer of our results in field conditions remain to be evaluated as other parameters could have influenced our results (cultivar used, no host-choice for aphids, growing conditions, substrate…). On the other hand, high-throughput sequencing (HTS) is increasingly used to identify viruses infecting cultivated and wild hosts [56–58]. Beyond virus detection, HTS technologies can also captures most of the genetic diversity of the viral population, as we have done by reconstructing viral haplotypes. In the future, implementing HTS in epidemiological survey programs, including on pooled samples as shown for Uganda cassava brown streak disease in Rwanda [29], would make it possible to identify the viral haplotypes and specific mutations present in the field, confirming hypotheses from experiments in controlled conditions and, in the long term, to deploy the most suitable plant genotypes for achieving a sustainable plant disease management.

## Materials and methods

### Virus and plant material

*Barley yellow dwarf virus* (BYDV; genus *Luteovirus*, family *Tombusviridae*) population used in this study was collected in 2013 (Pecq, Belgium) and maintained on six-row barley (*Hordeum hexastichum*) plants. A mass rearing of *Rhopalosiphum padi* aphids infected with the BYDV population was established on *H. hexastichum* plants in a controlled room (19–21°C, 16-h light/day). In 2017, fifteen months before the experiment, the BYDV population was transferred to and maintained on *Triticum aestivum* (unknown cultivar) using *R. padi*. This population was further used to perform the experimental evolution.

Four host species belonging to the *Poaceae* family were used to perform the experimental evolution: *Triticum aestivum* cv. Avatar, *Hordeum vulgare* cv. Laureate, *Hordeum hexastichum* cv. Smooth and *Avena sativa* cv. Evita. All the plants were grown under greenhouse conditions.

### Experimental evolution

At the start of the experiment, fourth-instar nymphs and adult apterous *R. padi* aphids, infected with both BYDV-PAV and BYDV-PAS (GenBank accessions OM046619 and OM046620), were placed on each host species. Fifteen plants per host species were used, with five aphids per plant. All the plants were inoculated four weeks after sowing. The five viruliferous aphids per plant were kept in a clip cage attached to the second leave of each plant. All aphids were manually removed after a four-days inoculation access period. Four weeks after inoculation, five virus-free *R. padi* aphids (fourth-instar nymphs and adult apterous) per plant were deposited in a clip cage set on the infected plants for a four-days acquisition access period. All virus-free aphids were reared on the same host species on which they were deposited during the experiment. Finally, the clip cages with the viruliferous aphids were attached on new plants of the same species to initiate a new passage. The virus was passed in the same host species for four passages of 4 weeks (Fig 1). At the end of a passage, each organ of each plant was homogeneously cut into several parts, half of it being kept to perform enzyme-linked immunosorbent assay (ELISA), and the other half being kept for HTS or back-up, according to the passage. All the samples were kept at -80°C. The experiment was performed in a climate-controlled growth room (19–21°C, 16-h light/day).

### Infection status of individual plants and extinction

The infectious status of each individual plant was assessed at each passage from plant A to plant B. Qualitative, double antibody sandwich enzyme-linked immunosorbent assay (DAS-ELISA) using the ELISA reagent set for *Barley Yellow Dwarf Virus*-PAV from Agdia (Elkhart, IN, USA). If the plant A was tested negative and, after 4 weeks, the plant B was also tested negative, the line was considered as extincted.

### Viral accumulation

Viral accumulation was assessed on each plant after the 1^st^ and 4^th^ passages. A quantitative double antibody sandwich enzyme-linked immunosorbent assay (DAS-ELISA) was performed using the same kit as above.). Polyclonal antibodies that recognize both BYDV-PAV and BYDV-PAS were used. DAS-ELISA was performed as described in [59]. Serial dilutions of infected plant samples were performed, and viral concentration in each plant was estimated relative to a standard control sample included on each ELISA plate. This allowed for the calculation of a relative concentration, indicating how many times higher or lower the viral concentration was in each plant compared to the control sample. After the grinding step, three and four replicates per plant were performed for the plants from the 1^st^ and 4^th^ passage, respectively. A qualitative ELISA was also performed on each plant after the 2^nd^ and 3^rd^ passages to check their infection status. Finally, a quantitative ELISA was performed to measure the viral accumulation of 4 viral populations (O1, O3 population P7, SB1 and SB3 population P11) on 3 hosts (wheat, oat and six-row barley) one month after inoculation, with 5 *R. padi* aphids per plant and four plants per treatment.

### Total RNA extraction and sequencing of the viral populations

Total RNA extraction was performed on one wheat plant infected with the initial BYDV population and on pools of plants from the 4^th^ passage. Plants from the 4^th^ passage were pooled together according to their viral accumulation, with two to five pools per host species. One oat plant (O1) and one six-row barley plant (SB1) were also extracted and sequenced alone because of their particularly high virus concentration (Fig 1). One gram of homogenized tissue from each plant was grounded in liquid nitrogen, the plants being pooled together during the grinding step. For the initial population, a different sampling protocol was followed: one gram of leaf tissue from 3 leaves was grounded in liquid nitrogen. Then, 120 mg of plant powder were used to perform total RNA extraction using 1.5 ml of TRIzol LS reagent (Thermo Fisher Scientific) per sample. After a 1-minute vortexing step and an incubation of 5 minutes at room temperature, 0.3 ml of chloroform were added to the samples. Samples were vortexed and incubated again for a 5-minutes period at room temperature, followed by a centrifugation at 4°C for 15 minutes at 13,000 rpm. The aqueous phase was collected and 0.75 ml of isopropanol were added. Samples were vortexed and incubated for 10-minutes at room temperature, followed by a centrifugation at 4°C for 20 minutes at 13,000 rpm. Then, the supernatant was discarded and the pellet washed with 75% ethanol, followed by a centrifugation at 4°C for 5 minutes at 9,000 rpm. The supernatant was discarded and the pellet dried for 10 minutes. The pellet was resuspended in 100 μl of water and an equal volume of phenol:chloroform:isoamyl alcohol (25:24:1) was added to each sample, followed by a centrifugation at 4°C for 2 minutes at 13,000 rpm. The supernatant was collected and 10% volume of sodium acetate 3M pH 5.2 as well as 80% volume of isopropanol were added. Samples were kept at -20°C for an overnight precipitation. After a centrifugation step at 4°C for 20 minutes at 13,000 rpm, the supernatant was discarded and 1 ml of 75% ethanol was added to wash the pellet. After another centrifugation step at 4°C for 5 minutes at 9,000 rpm, the supernatant was discarded and the pellet dried for 10 minutes at room temperature. Finally, the pellet was resuspended in 100 μl of nuclease-free water.

Samples were then processed by the GIGA-Genomics platform of Liège University (Liege, Belgium). A ribo-depletion procedure was first carried out using the RiboMinus Plant Kit for RNA-Seq (Thermo Fisher Scientific). Then, a total RNA library was constructed with the TruSeq Stranded mRNA kit (Illumina). Finally, samples were sequenced using the Illumina NextSeq 500 system (2×150 nucleotides).

### Bioinformatic analyses

Bioinformatic analyses were performed using Geneious software (version 10.1.3, Biomatters). A total of 12,701,158 illumina paired-reads were obtained for the initial BYDV population. The number of paired-reads ranged between 9,845,404 and 11,806,570 reads for the fourteen pools of plants infected with the final BYDV populations. The obtained reads were paired, all the reads being paired with each other for all the samples, and merged. Then, a quality trimming step was performed on both the merged and unmerged paired-reads using the BBDuk plugin (version 38.84), with a minimum quality score of 30 and a minimum read length of 20 bp. Finally, dedupe was applied to remove duplicate reads on both the merged and ummerged trimmed paired reads. The reads obtained were all used for the next steps of the analysis.

*De novo* assembly was performed on the initial BYDV population using SPAdes assembler (version 3.13.0) [60]. Contig annotation was performed using both blastn and tblastx programs against the NCBI nucleotide and viral RefSeq databases (release versions 231 and 92), respectively. After the identification of the contigs assigned to either BYDV-PAV or BYDV-PAV, another *de novo* assembly step was carried out with SPAdes using the BYDV contigs assigned to each species and two complete BYDV genomes were obtained.

Reads from the final BYDV populations were mapped at the same time on the BYDV-PAV and the BYDV-PAS references obtained previously. When multiple best matches occurred, the reads were discarded. A complete coverage of the references was obtained for all the samples, except when one of the BYDV species was not present in the sample. The mappings obtained were used to perform SNP calling. Filtering of the SNPs was carried out according to the following criteria: (i) a minimum coverage of 100x, (ii) a minimum Strand-Bias > 65% p-value of 0.0005, the SNPs with a smaller strand bias p-value being excluded, and (iii) a minimum variant frequency of 1%. The Maximum-likelihood (ML) trees of the haplotypes were generated with Geneious software (version 10.1.5, Biomatters) using the PhyML plugin (version 3.3.20180621) under the HKY85 model of sequence evolution.

### FST measures and haplotype reconstruction

Pairwise FST between all final BYDV populations were calculated separately for BYDV-PAS and BYDV-PAV populations. FST measures were obtained using Popoolation2 (version 1.201) [61], with a window size equal to the viral genome size (5,570 nt for BYDV-PAV and 5,577 nt for BYDV-PAS), a step size of 1, a minimum covered fraction of 0.1, a minimum coverage of 50x and a maximum coverage of 200,000x. Both the classical approach [62] and the approach adapted to digital data [63] were applied and gave very similar results. Therefore, only the results obtained with the classical approach are presented.

Haplotype reconstruction was performed on two genome regions for BYDV-PAS (850-1,566 nt and 3,630-5,172 nt, on BYDV-PAS, GenBank accession OM046619) and one genome region for BYDV-PAV (4,000-5,111 nt, on BYDV-PAV, GenBank accession OM046620). These regions were selected because of their high SNP density, two SNPs being no more distant than 150 nt, which increases the odds of finding at least two SNPs on the same read and makes the haplotype reconstruction easier and more robust. Pre-processed reads were mapped on these regions as explained before in order to obtain Sequence Alignment Map (SAM) files for each sample. Clique SNV [53] was applied to perform reconstruction of viral haplotypes, with a minimum haplotype frequency of 0.01.

### Statistical analysis

All statistical analyses were performed using the R software (http://www.r-project.org/). A Wilcoxon test was used to compare viral accumulation mean values between initial (1^st^ passage) and final (4^th^ passage) viral populations, with the wilcox.test function implemented in the package stats. A Dunnett test was used to compare the viral accumulation between every final BYDV population and the initial BYDV population in each host species using the nparcomp package. Principal component analyses were carried out using the ade4 and factoextra packages. The dendrograms were calculated using the pvclust package. The statistical support of the clusters was assessed using multi-scale bootstrapping, with 10,000 bootstrap replicates. For each cluster, an approximately unbiased (AU) p-value was obtained and used to evaluate the uncertainty of the hierarchical clustering. For AU p-values > 0.95, the cluster was considered “valid” (*i.e.,* strongly supported by the data) at a significance level of 0.05.

## Supporting information captions

**S1 Figure: Principal Component Analysis.** The principal component analysis shows the first and second dimensions obtained using all the detected SNPs as variables for BYDV-PAS (A) and BYDV-PAV (B). The proportion of BYDV-PAS in each viral pooled population is also shown as a function of position along PCA dimension 1 and viral fitness (C). Colour of the points indicates the host species in which the BYDV pooled populations evolved: wheat (green), oat (grey), two-row barley (blue) or six-row barley (red). Names of the viral populations are indicated (W1 to W3 for wheat; O1 to O3 for oat; TB1 to TB3 for two-row barley; SB1 to SB5 for six-row barley). In panel C, circle size is proportional to fitness (virus accumulation in the plant) level. In the PCA analysis (panels A and B), the fitness of the analysed pooled populations was not considered. In contrast, panel C incorporates fitness data; however, SB1, which exhibited an exceptionally high fitness level, was marked as a square which whose size is not proportional to the virus fitness.

**S1 Table: Infection status of the plants after each passage.** Presence of BYDV in each plant was controlled after every passage with either a qualitative or a quantitative ELISA. A ‘1’ means that the plant was infected with BYDV and a ‘0’ means that the plant was not infected with BYDV.

**S2 Table: Pairwise distance matrix for the alignment of NCBI BYDV references and the two BYDV genomes from this study.** The matrices have been obtained after the multiple alignment of 113 BYDV nucleotide sequences from NCBI and the two BYDV genomes generated in this study (GenBank accessions OM046619 and OM046620). BYDV genomes from this study are highlighted in red.

**S3 Table: Number of reads mapped against BYDV-PAS and BYDV-PAV genomes as well as the average read depth for each viral pooled population.**

**S4 Table: Description of the haplotypes reconstructed for each viral population.** Haplotype reconstruction was performed on three different genomic regions, two for BYDV-PAS and one for BYDV-PAV. The number of reconstructed haplotypes for each region, their frequencies in the viral population as well as their group are given.

## Notes

### Competing Interest Statement

The authors have declared no competing interest.

### Summary of Updates

Yellow dwarf viruses are damaging viruses infecting cereals. They are able to infect a wide range of host plants belonging to the Poaceae family. The ban of neonicotinoids in Europe has resulted in an increasing disease incidence and triggered the need to better understand their emergence and spread. The ability of a Barley yellow dwarf virus (BYDV) population to adapt to different hosts has never been studied. We performed an experimental evolution of two BYDV species (BYDV-PAS and BYDV-PAV) to study their adaptation to four Poaceae species (wheat, oat, two-row barley, and six-row barley). After four months of evolution (4 passages from plant to plant), the replicative fitness of the final viral populations was estimated, and the complete viral genomes were sequenced by high-throughput sequencing in pools of BYDV populations. Wildly divergent evolutionary trajectories were obtained, with stable or increased fitness, up to extinctions of viral populations within and among plant species. To understand these results, the composition of viral populations was analysed in detail using single nucleotide polymorphism (SNP) calling, clustering, and haplotype reconstruction methods. Interestingly, adaptation to oat and barley was mainly explained by a combination of BYDV-PAV haplotypes showing specific mutations. In contrast, adaptation to wheat was mainly explained by a combination of BYDV-PAS haplotypes harbouring specific mutations. Moreover, these local adaptations were associated to an adaptation cost in other hosts for some viral populations. The presence of adaptation costs in controlled but realistic conditions opens the door for evaluating practices such as crop mixtures or rotations on fields, as a means to mitigate the impact of BYDV.

### Funding

- Special Research Funds (FSR) of Liège University

