## Supplementary figures and images for "Host-specific adaptation and fitness trade-off of Barley Yellow Dwarf Viruses suggested by experimental evolution through aphid inoculation on multiple Poaceae species"

### Figure S1

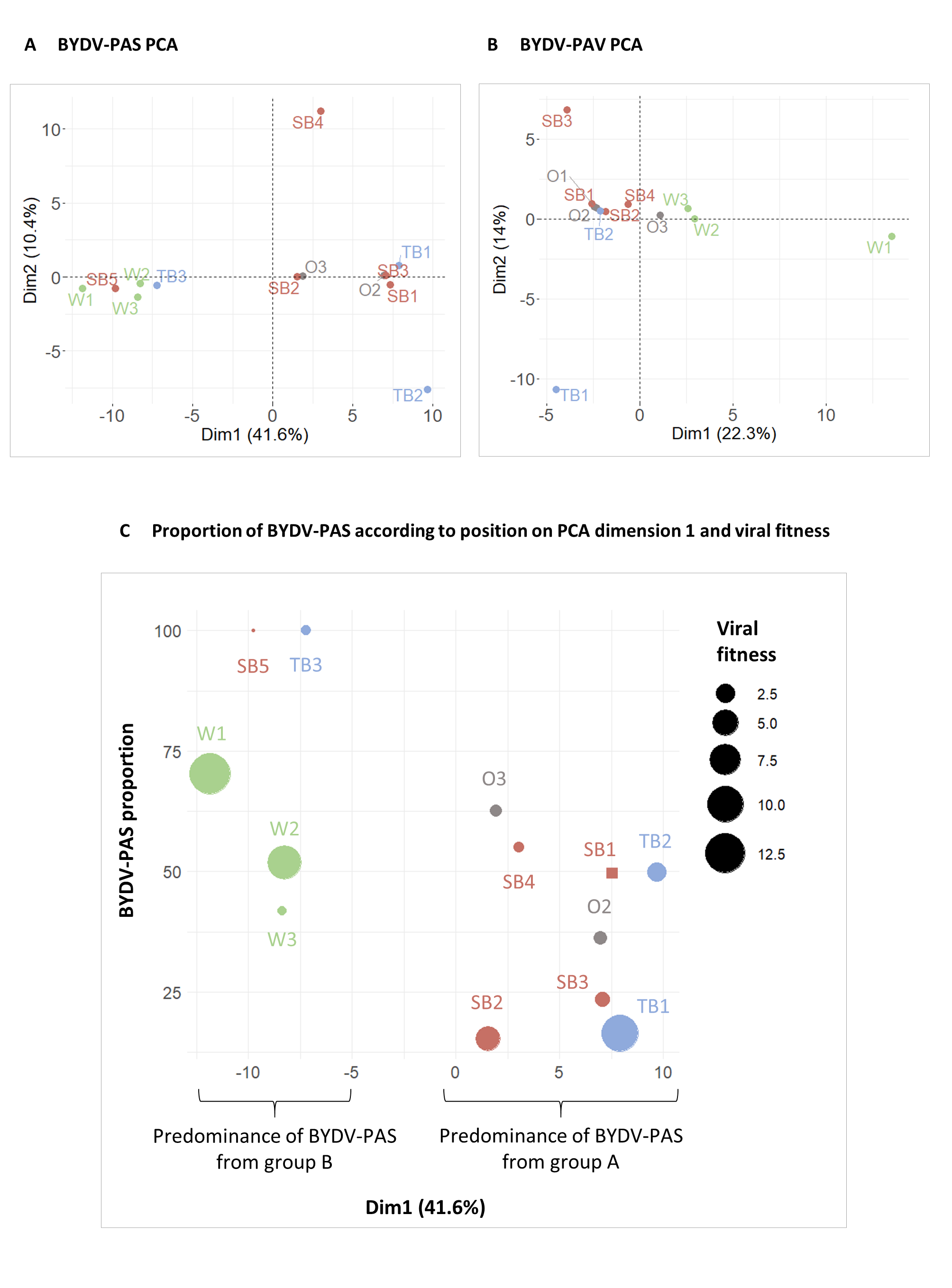
